# SweepCluster: A SNP clustering tool for detecting gene-specific sweeps in prokaryotes

**DOI:** 10.1101/2021.03.12.435060

**Authors:** Junhui Qiu, Qi Zhou, Weicai Ye, Qianjun Chen, Yun-Juan Bao

## Abstract

**Background:** The gene-specific sweep is a selection process where an advantageous mutation along with the nearby neutral sites in a gene region increases the frequency in the population. It has been demonstrated to play important roles in ecological differentiation or phenotypic divergence in microbial populations. Therefore, identifying gene-specific sweeps in microorganisms will not only provide insights into the evolutionary mechanisms, but also unravel potential genetic markers associated with biological phenotypes. However, current methods were mainly developed for detecting selective sweeps in eukaryotic data of sparse genotypes and are not readily applicable to prokaryotic data. Furthermore, some challenges have not been sufficiently addressed by the methods, such as the low spatial resolution of sweep regions and lack of consideration of the spatial distribution of mutations.

**Results:** We proposed a novel gene-centric and spatial-aware approach for identifying gene-specific sweeps in prokaryotes and implemented it in a python tool SweepCluster. Our method searches for gene regions with a high level of spatial clustering of pre-selected polymorphisms in genotype datasets assuming a null distribution model of neutral selection. The pre-selection of polymorphisms is based on their genetic signatures, such as elevated population subdivision, excessive linkage disequilibrium, or significant phenotype association. Performance evaluation using simulation data showed that the accuracy and sensitivity of the clustering algorithm in SweepCluster is above 90%. The application of SweepCluster in two real datasets from the bacteria *Streptococcus pyogenes* and *Streptococcus suis* showed that the impact of pre-selection was dramatic and significantly reduced the uninformative signals. We validated our method using the genotype data from *Vibrio cyclitrophicus*, the only available dataset of gene-specific sweeps in bacteria, and obtained a concordance rate of 78%. We noted that the concordance rate could be underestimated due to distinct reference genomes and clustering strategies. The application to the human genotype datasets showed that SweepCluster is also applicable to eukaryotic data and recovered the known sweep regions in a wide dynamic range of pre-selection parameters.

**Conclusions:** SweepCluster is applicable to a broad category of datasets. It will be valuable for detecting gene-specific sweeps in diverse genotypic data and provide novel insights on adaptive evolution.

## Background

A selective sweep is a process where a beneficial allelic change sweeps through the population and becomes fixed in a specific population, and the nearby sites in linkage disequilibrium will hitchhike together and also become fixed (1, 2). Those sweep regions containing beneficial alleles could possibly be introduced by recombination and rise to high frequency rapidly in the population under positive selection. If the increase in frequency is recent or fast relative to other recombination events, the mutation profile in the sweep regions across the population will be maintained without being interrupted. Finally, the process will imprint genetic signatures in the population genomes, leading to lowered within-population genetic diversity, increased between-population differentiation, and/or high linkage disequilibrium (3-5). When such selective sweeps only occur at specific gene regions under selection without affecting the genome-wide diversity, they are described as gene-specific sweep (6).

Recently, the gene-specific sweep has been demonstrated to play important roles in adaptive evolution in microbial populations, such as ecological differentiation in Prochlorococcus (7) and Synechococcus (8), speciation in marine bacterium *Vibrio cyclitrophicus* (*V. cyclitrophicus*) (3, 9), and phenotypic divergence in human adapted pathogen *Streptococcus pyogenes* (*S. pyogenes*) (10). The observation of the gene-specific sweeps in those scenarios in both environmental organisms and host pathogens suggests that the gene-specific sweep may represent one of the general mechanisms underlying adaptive evolution of microorganisms. Therefore, identifying the gene-specific sweep on the genome-wide scale will not only provide insights into the evolutionary mechanisms shaping the genetic diversity, but also help to unravel potential genetic markers associated with ecological adaptation or phenotypic differentiation.

An array of methods have been proposed to identify the gene-specific sweeps and are generally fall into three categories based on the genetic signatures being captured, *i*.*e*., (1) composite likelihood ratio (CLR) tests of the marginal likelihood of the allele frequency spectrum under a model with selective sweeps in comparison with that under a model of selective neutrality (11-13) (Kim and Stephan-2002, Nielsen-2005, Huber-2016); (2) comparison of the distribution of population subdivision or linkage disequilibrium in a region under positive selection with that of a neutral background (14, 15) (Akey-2002, Kim-Nielsen-2004); (3) haplotype-based approaches for detecting elevated haplotype homozygosity in a locus around the selected site in comparison with that under a neutral model (16-20) (Sabeti-2002, Voight-2006, Ferrer-Admetlla-2014, Harris-2018, Harris-2020). Those methods have demonstrated the power for detecting genetic signatures of selective sweep in numerous cases.

However, those methods were mainly developed for detecting selective sweeps in eukaryotic data and are not readily applicable to prokaryotic data, such as the haplotype-based approaches (21). In addition, some challenges have not yet been sufficiently addressed by the currently available methods. For example, the gene-centric concept of the gene-specific sweep has not been taken into account leading to a low spatial resolution of sweep regions; the spatial distribution properties of the mutated sites within the sweep regions have not been fully considered.

In this study, we propose a new gene-centric approach for identifying the gene-specific sweeps in prokaryotes, which search for regions with a higher level of spatial clustering of single nucleotide polymorphisms (SNPs) assuming a null distribution model of SNPs under neutral selection. The clustering applies to the SNP subsets of specific interests, which can be selected based on the genetic signatures of sweep regions, such as elevated population subdivision, reduced within-population diversity, excessive linkage disequilibrium, or significant phenotype association. Our approach is different from the previous methods in that: (1) it applies the gene-centric concept by considering the gene-specific location of SNPs; (2) it takes advantages of the spatial distribution properties of SNPs in the sweep region; (3) the clustering is performed on pre-selected target SNPs with specific genetic properties, thus minimizing the influences from uninformative SNPs. We offer it as an open-source tool “SweepCluster” and it is freely accessible at github: https://github.com/BaoCodeLab/SweepCluster.

## Methods

### Pre-selection of SNPs

The pre-selection of SNPs could be based on elevated population differentiation *F*st, extended linkage disequilibrium LD, or phenotypic association. However, the determination could also depend on the data property and study purposes. For instance, if the positive selection acting on disease markers is of interest, the screening of SNPs with significant association with disease phenotypes using robust genome-wide association analysis is preferred. In the real and simulated datasets in this study, we selected the SNPs associated with phenotypic divergence or population differentiation.

### Overview of the clustering approach

The SNP clustering algorithm employs a gene-centric concept to mimic the biological process of introducing gene-specific sweeps. In the gene-specific sweep model, non-synonymous SNPs (the SNPs causing amino acid alterations) or upstream regulatory SNPs (the SNPs in the regulatory regions) are more likely to be under positive selection than synonymous SNPs (the SNPs without causing amino acid alterations) or inter-genic SNPs, and the selected non-synonymous SNPs along with the nearby synonymous/inter-genic SNPs are introduced simultaneously in a single event. For a recent sweep event, the selected SNPs and the hitchhiking SNPs are tightly clustered in specific gene regions without severely ruined by other recombination events. Based on the gene-specific sweep model, our clustering strategy is illustrated in Figure 1 and described previously (10). Briefly, a non-synonymous or upstream regulatory SNP is randomly chosen in a specific gene/operon and serves as an anchor for an initial cluster. The initial cluster is then extended progressively by scanning and merging the neighboring SNPs or clusters. If the total span is shorter than the specified sweep length, then the surrounding SNPs or clusters are merged. Otherwise, the initial cluster is extended by merging the neighboring SNPs or clusters which minimize the normalized root-mean-square of inter-SNP distances (NRMSD):

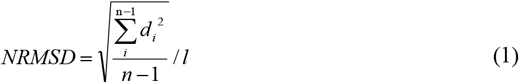

where *d*_*i*_ is the *i*^th^ inter-SNP distance, *n* is the total number of the SNPs in the target cluster and *l* is the maximum spanning range of the SNPs in the target cluster.

**Fig. 1.**
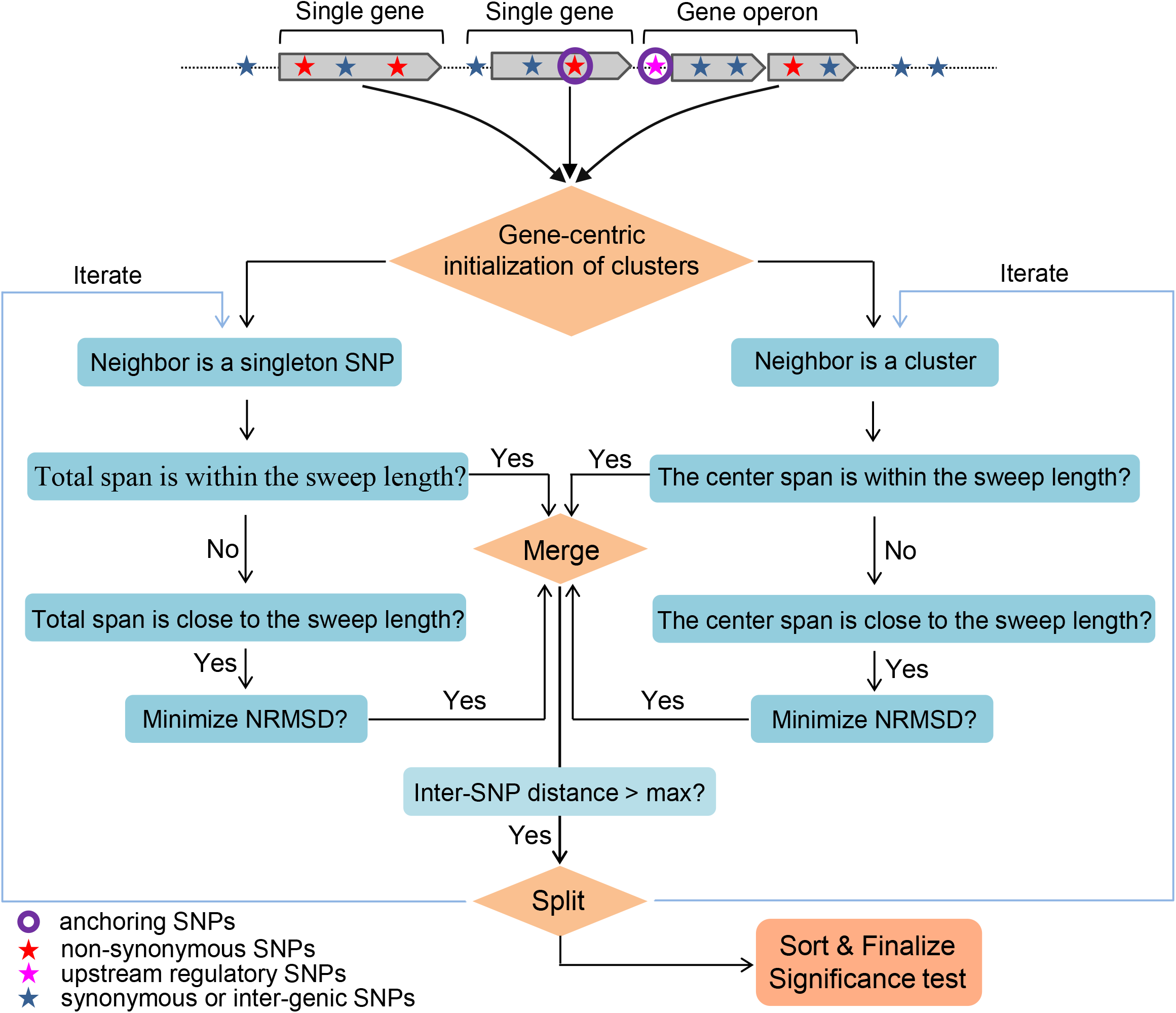
Outline of the clustering procedures of SweepCluster. A non-synonymous or upstream regulatory SNP is randomly chosen in each gene/operon and serves as an anchoring SNP for an initial cluster. The initial cluster is then extended by scanning and merging the neighboring SNPs or clusters. If the total span is shorter than the specified sweep length, then the surrounding SNPs or clusters are merged. Otherwise, the initial cluster is extended by merging the neighboring SNPs or clusters such that the normalized root-mean-square of inter-SNP distances (NRMSD) is minimized. All clusters after merging are re-examined and split if any inter-SNP distance within the cluster is longer than a given inter-SNP distance threshold (max_dist).

Following merging, all clusters are re-examined and split if any inter-SNP distance within the cluster is longer than a given distance threshold. The distance threshold can be determined based on the genome-wide average inter-SNP distance. Under the null neutral model, the SNPs are independently and randomly distributed across the genome, and the significance of a cluster with *m* distinct SNPs spanning a length of *l* can be evaluated using the gamma distribution with the average mutation rate μ as the rate parameter (22):

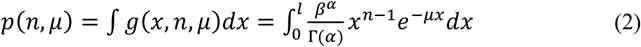

The average mutation rate across the genome can be calculated as: *μ* = *n*/*s*, where *n* is the total number of SNPs in the genome and *s* is the length of the genome.

### Evaluation of the performance of the clustering method

We evaluated the performance of the clustering using four metrics, *i*.*e*., CPU time, memory usage, accuracy and sensitivity based on simulation datasets. The evaluation of CPU time and memory usage was performed using real datasets with varying data size. The assessment of accuracy and sensitivity was conducted based on simulation datasets (see below). The accuracy is defined as the proportion of correctly assigned SNPs among the total SNPs. The sensitivity is defined as the proportion of detected clusters containing at least 90% of the SNPs correctly assigned. The mapping between detected clusters and expected clusters was determined based on reciprocal maximum overlapping between the two sets of clusters.

### Simulation datasets

The simulation datasets for assessing the accuracy and sensitivity of the clustering algorithm were generated based on the genome and annotation of the bacterial strain *S. pyogenes* AP53, which was annotated and studied by us previously (23). The SNPs were artificially generated independently and randomly on the genome based on the Poisson process of a given mutation rate (the average mutation rate of *S. pyogenes*). SNP clusters were then created by taking the following procedures to satisfy the pre-defined threshold of sweep length (sweep_lg) and maximum inter-SNP distance (max_dist): (i) roughly a half of the SNPs in each gene region were assigned non-synonymous; (ii) removing the SNPs in the gene regions longer than sweep_lg+50; (iii) if the spanning length of the neighboring genes is longer than sweep_lg+50 and the inter-genic distance is greater than max_dist, then remove the downstream gene to create a larger inter-genic distance.

### Real dataset of *S. pyogenes* genotypes

We used the real genomic datasets from two bacterial species *S. pyogenes* and *S. suis* to assess the effects of the procedure of SNP pre-selection. The reason to choose the two species is that they are known to have a high level of genomic varibility and a high density of genotypes, which facilitate the manifestation of influences of SNP pre-selection (24,25). *S. pyogenes* is a common human pathogenic bacterium causing diverse disease phenotypes, such as pharyngitis, skin infection, necrotizing fasciitis, and acute rheumatic fever. Previous studies have shown that the alleles in the gene regions of *S. pyogenes* exhibit phenotype-dependent changes, thus providing an excellent dataset for selecting SNPs associated with phenotype differentiation (10, 26). The genomic sequences of *S. pyogenes* were downloaded from NCBI Genbank database (ftp://ftp.ncbi.nlm.nih.gov). A total of 46 genomes were chosen for this study with balanced distribution of phenotypes based on the known phenotypic information (10). The core genome is defined as the regions encoded by all studied genomes and was determined by aligning the shredded genomes against the reference strain AP53 (CP013672). Finally, the core genome contains 69,171 segregating sites mutated in at least one of the genomes and were concatenated for downstream analysis. Both the whole set of SNPs at all segregating loci and a subset of selected SNPs associated with the phenotype of acute rheumatic fever were used for inferring sweep regions using SweepCluster. The SNPs associated with the disease phenotype were identified using the Chi-squared test. The parameters used for SweepCluster are “-sweep_lg 1781 -max_dist 1100 -min_num 2” and the clustering significance was evaluated using the function “Pval” with the parameter of mutation rate “-rate 0.0362”. The linkage disequilibrium analysis of the SNPs was performed using Haploview (27).

### Real dataset of *Streptococcus suis* (*S. suis*) genotypes

*S. suis* is a swine pathogen that colonizes pigs asymptomatically but can also causes severe clinical diseases in pigs such as respiratory infection, septicemia, and meningitis. *S. suis* can be classified to 29 distinct serotypes forming complex population structures (28). Previous phylogenetic study showed that many serotypes exist in multiple subpopulations and each subpopulation may contain multiple serotypes (25). The complexity has been associated with extensive genetic recombination and genomic shuffling among and between populations. Therefore, it will be interesting to investigate the occurrence of selective sweeps among subpopulations in the highly recombining genome of *S. suis*.

A total of 1,197 genomic sequences of *S. suis* strains were downloaded from the NCBI Genbank database (ftp://ftp.ncbi.nlm.nih.gov). We removed the redundancy among the genomes to reduce the data size by grouping them based on the submission institutions and selecting the most distant genomes within each group based on the phylogenetic structures built by SplitsTree (29) (Figuure S1A). The selected genomes were further filtered based on their phylogenetic distance. The final dataset comprises 208 non-redundant genomes (Figure S1B) and gives rise to a total of 236,860 segregating mutation sites with BM407 as the reference (FM252033). The core genome was identified using the same procedures as that for *S. pyogenes*. The inference of sweep regions using SweepCluster was performed respectively for all segregating SNPs and for those associated with differentiation of two subpopulations (branch-1 and branch-2 in Figure S1C). The SNPs associated with population differentiation was identified using the Chi-squared test. The parameters used for SweepCluster and significance evaluation are “-sweep_lg 2000 -max_dist 2000-min_num 2” and “-rate 0.1077”, respectively.

### Real dataset of *V. cyclitrophicus* genotypes

*V. cyclitrophicus* is a gram-negative bacterium inhabiting seawater. Previous studies reported ecological differentiation of the *V. cyclitrophicus* population associated with gene-specific sweeps (9). The authors sequenced 20 strains of *V. cyclitrophicus*, which are divided into two phenotypic groups (S strains and L strains) according to their ecological partition. They showed that the partition is associated with the ecoSNPs, *i*.*e*., the dimorphic nucleotide positions with one allele present in all S strains and the other allele in all L strains. The authors then classified the ecoSNPs into 11 clusters and demonstrated the evidences of gene-specific sweeps in causing the ecoSNPs. This is the only available study of SNP clusters under gene-specific sweeps in bacteria. We used this dataset for benchmarking of our clustering method.

We downloaded the genomic sequences of the 20 strains from NCBI Genbank database (ftp://ftp.ncbi.nlm.nih.gov) and aligned them to a reference strain with complete genome assembly (ECSMB14105) to derive the segregating SNPs of 139,066 and the phylogenetic structure (Figure S2). The ecoSNPs were obtained using the same definition as that in the reference (9). The ecoSNPs were then subject to cluster detection using SweepCluster with the parameters “-sweep_lg 8000-max_dist 5000 -min_num 2” and “-rate 0.000111”.

### Empirical datasets of human genotypes

We employed the genotype datasets from the human 1000 Genomes project (30) to evaluate the ability of SweepCluster of identifying selective sweeps in eukaryotic data. We chose the 1000 Genomes datasets because they have been extensively used in previous studies of selective sweeps and a handful of gene loci have been well-characterized to be under selective sweep in specific subpopulations. We extracted the genotype data from three subpopulations, *i*.*e*., EUR (Europeans), AFR (Africans) and EAS (East Asians), and selected the mutation sites associated with pairwise population differentiation *F*st. The calculation of *F*st was based on Hudson’s estimator in the transformed formula (31):

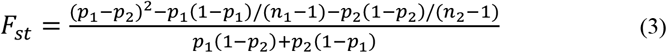

where *n*_1_/*n*_2_ is the subpopulation size and *p*_1_/*p*_2_ is the allele frequency for the two paired populations. Distinct subsets of SNPs were selected using a series of *F*st thresholds (0.7, 0.65, 0.60, 0.55, 0.50, 0.45, 0.43, and 0.4) for inferring sweep regions to evaluate the robustness of SweepCluster in eukaryotic data. The parameters used for SweepCluster are: “-sweep_lg 200000 –max_dist 40000 –min_num 2”. The sweep regions and SNPs were annotated based on the genome build hg19 using ANNOVAR (32).

### Optimization of the parameters

We carried out the parameter simulation of sweep lengths by calculating the number of sweep regions inferred by SweepCluster for varying values of sweep lengths in the range 300-10,000 bp. The relationship between the number of sweep regions versus sweep length was approximated using non-linear fitting implemented in generalized additive models in the R package “mgcv”. The optimal estimation of the sweep length is calculated based on the maximum curvature in the fitting curves with the curvature calculated with the following formula:

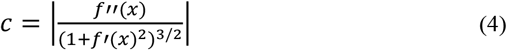

where *f*′(*x*) and *f*′′(*x*) are the first-order and second-order derivative of the fitting curves, respectively. We have provided in the package a shell script “sweep_lg_simulation.sh” for automatic optimization of the sweep length for any particular genotype dataset. The parallel acceleration was implemented in the script for fine-grained parameter searching.

## Results

### Overview of SweepCluster

The package SweepCluster performs four major functions. (1) Density: calculates the SNP density using a window-scanning method in a specific genomic region or in the genome-wide scale; (2) Cluster: executes the core functionality of the package, *i*.*e*., gene-centric SNP clustering; (3) Pval: estimates the statistical significance of each SNP cluster based on a null gamma distribution of SNPs; (4) a driver script “sweep_lg_simulation.sh” for parameter optimization.

### Computing performance

The computing performance of SweepCluster was evaluated using multiple real datasets with varying number of SNPs (designated as *N*). The memory usage of SweepCluster increases linearly with *N* and is fairly low even for the maximum datasets of 200,000 SNPs at about 260 megabytes (MB) (Figure 2A). The CPU time consumption of SweepCluster is on the scale O(*N*^2^) at the initial stage and then becomes nearly linear O(*N*) when *N* > 140,000 (Figure 2B). It is because the CPU time is governed by optimizing the boundary SNPs when *N* is small, but becomes governed by clustering the inner SNPs for large *N*s, at which the ratio of boundary SNPs rapidly declines. Considering the linear increment of memory usage and CPU time, and the downsized genotype datasets upon pre-selection, we anticipate that the computing resources will not be limiting factors for larger datasets. In the meanwhile, it should also be noted that the computing performance also depends on the applied parameters (such as the sweep length) and the genotype data properties (such as the proportion of the boundary SNPs).

**Fig. 2.**
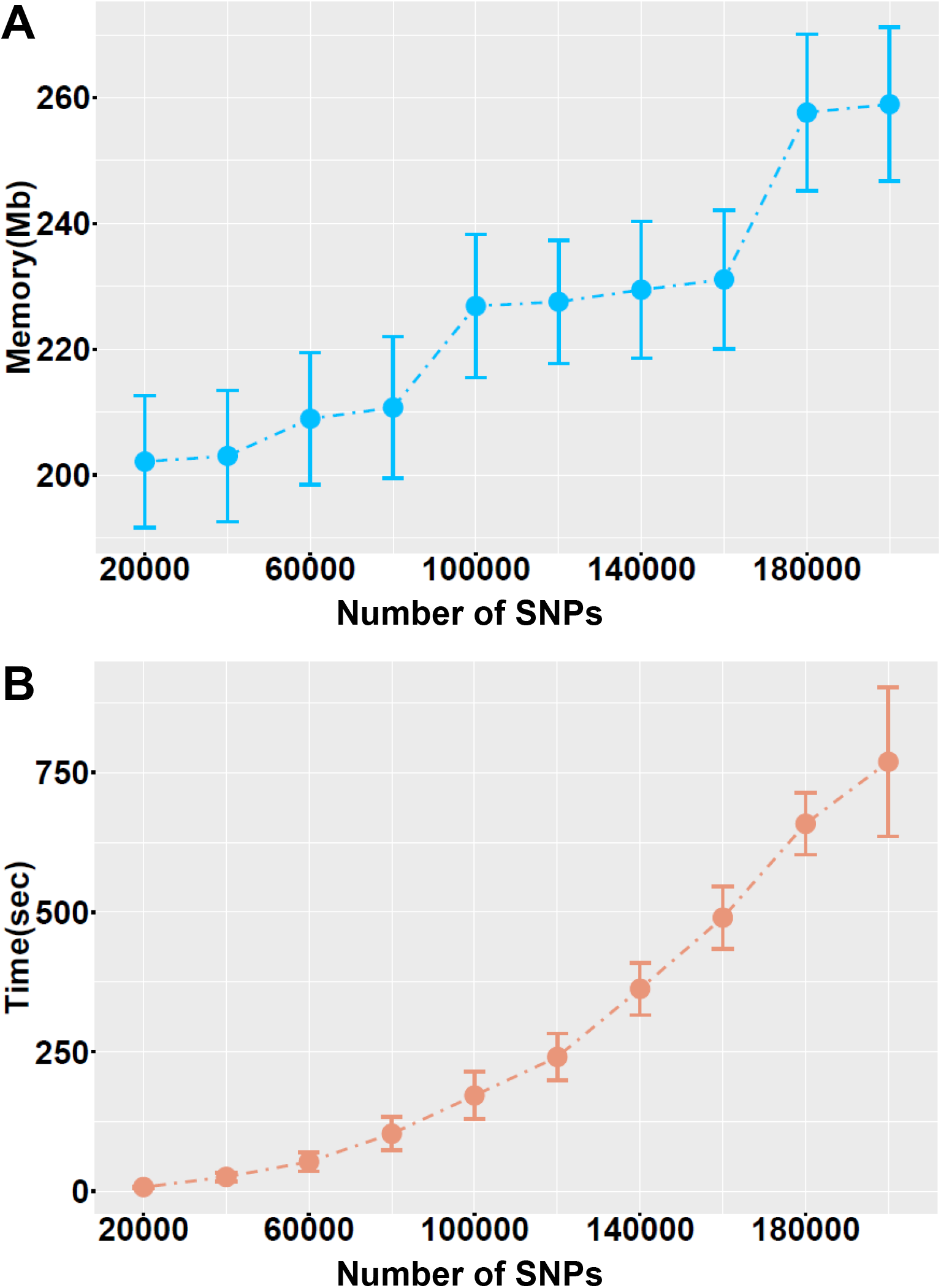
Memory usage (A) and CPU time (B) of SweepCluster for varying numbers of SNPs. The datasets for evaluation were obtained by subsetting the genotype dataset of *S. suis*.

### Performance of accuracy and sensitivity

We evaluated the performance of the clustering algorithm in SweepCluster in terms of accuracy and sensitivity using artificially generated simulation datasets with the SNP distributions satisfying specific combinations of sweep lengths (sweep_lg) and maximum inter-SNP distances (max_dist). The performance of SweepCluster was compared with that of DBSCAN, a general-purpose and commonly used spatial clustering algorithm without considering any trait information of the data (33). The comparison showed that the performance of both algorithms as a function of maximum inter-SNP distances is highly similar, where the accuracy and sensitivity quickly approaches optimum when the maximum inter-SNP distance increases to roughly 200 bp, close to the average inter-SNP distance in the gene regions in the simulation datasets (Figure 3A,C). Interestingly, the performance of SweepCluster and DBSCAN as a function of sweep lengths differs (Figure 3B,D). DBSCAN is not influenced significantly by the sweep length and performs nearly equally well for a broad range of sweep lengths. However, the performance of SweepCluster is dependent on the sweep length. It gradually improves with increasing sweep lengths and achieves optimal results at around 800-1000 bp, coincident with the average gene length of our simulation datasets. The dependence of the performance of SweepCluster on the sweep length is a manifestation of the gene-aware concept of the design of the clustering method in SweepCluster. In biological contexts, the general-purpose clustering methods, such as DBSCAN may generate clusters unrelated with selective sweeps.

**Fig. 3.**
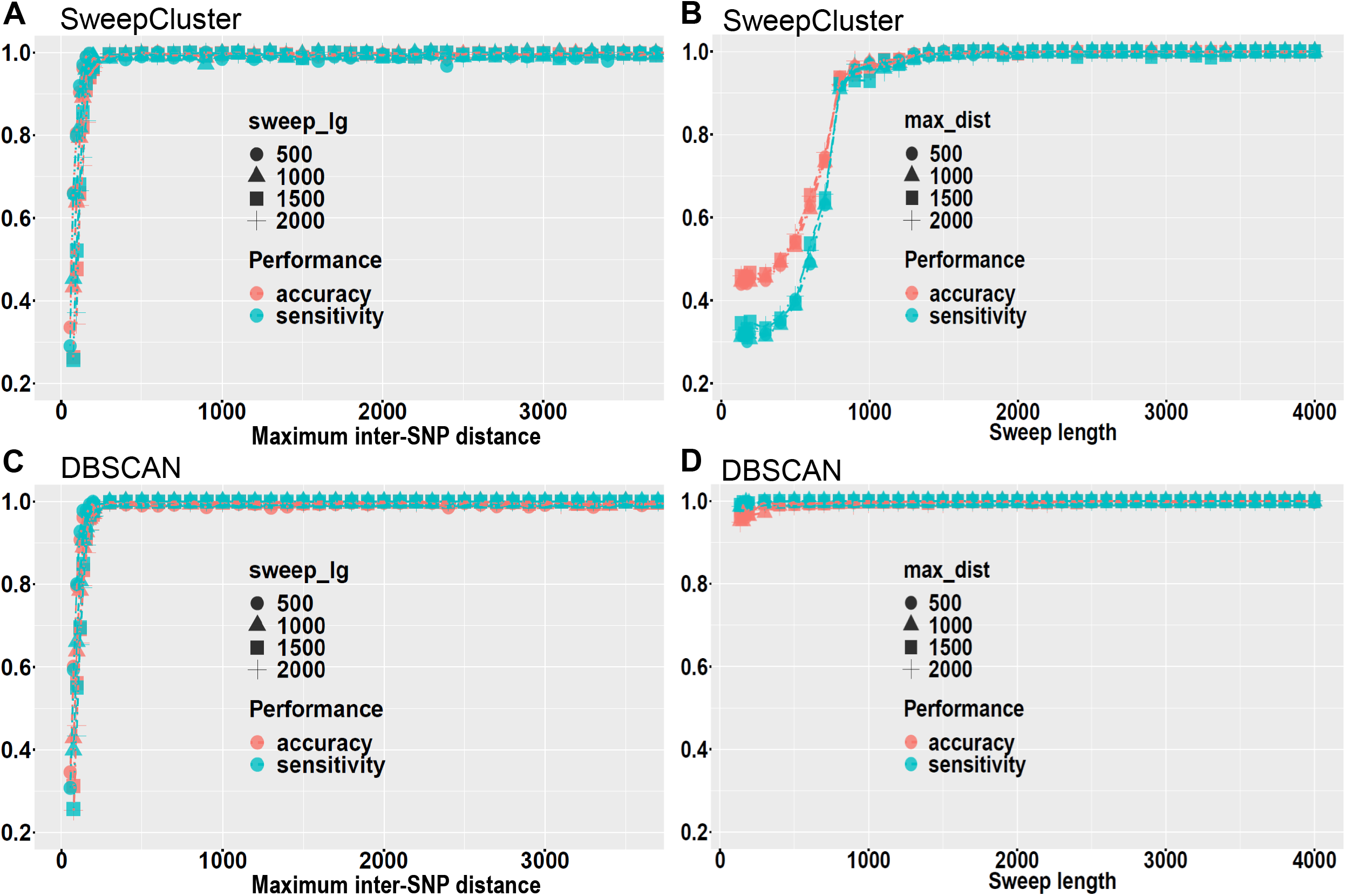
The accuracy and sensitivity of the clustering algorithm in SweepCluster in comparison with DBSCAN. The accuracy and sensitivity were calculated for a series of values of sweep lengths or maximum inter-SNP distances.

### Efficacy of SNP pre-selection in real datasets of *S. pyogenes and S. suis*

We test the efficacy of the procedure of SNP pre-selection prior to clustering by employing real datasets from two bacterial species, *S. pyogenes* and *S. suis* of dense genotypes.

For the datasets of *S. pyogenes*, a total of 69,171 core SNPs were obtained across 46 representative strains and selection of SNPs based on phenotypic association with the disease acute rheumatic fever reduced the number of SNPs to 1,631 (**Additional file 1: Table S1, S2 and S3**). SweepCluster was subsequently applied to the two SNP datasets and identified 215 and 131 significant clusters (*p*-value 0.05), respectively (Figure S3, **Additional file 1**: Table S4 and S5). We then used linkage disequilibrium (LD) between SNPs within the clusters as a proxy to examine the effect of pre-selection. A snapshot of the comparison of the LD patterns before and after pre-selection is shown in Figure 4A,B. The average LD within clusters was significantly increased after performing SNP pre-selection (*p*-value < 2.2 × 10^−16^), indicating the significant effect of pre-selection on diminishing the spurious signals in inferring sweep regions (Figure 4C).

**Fig. 4.**
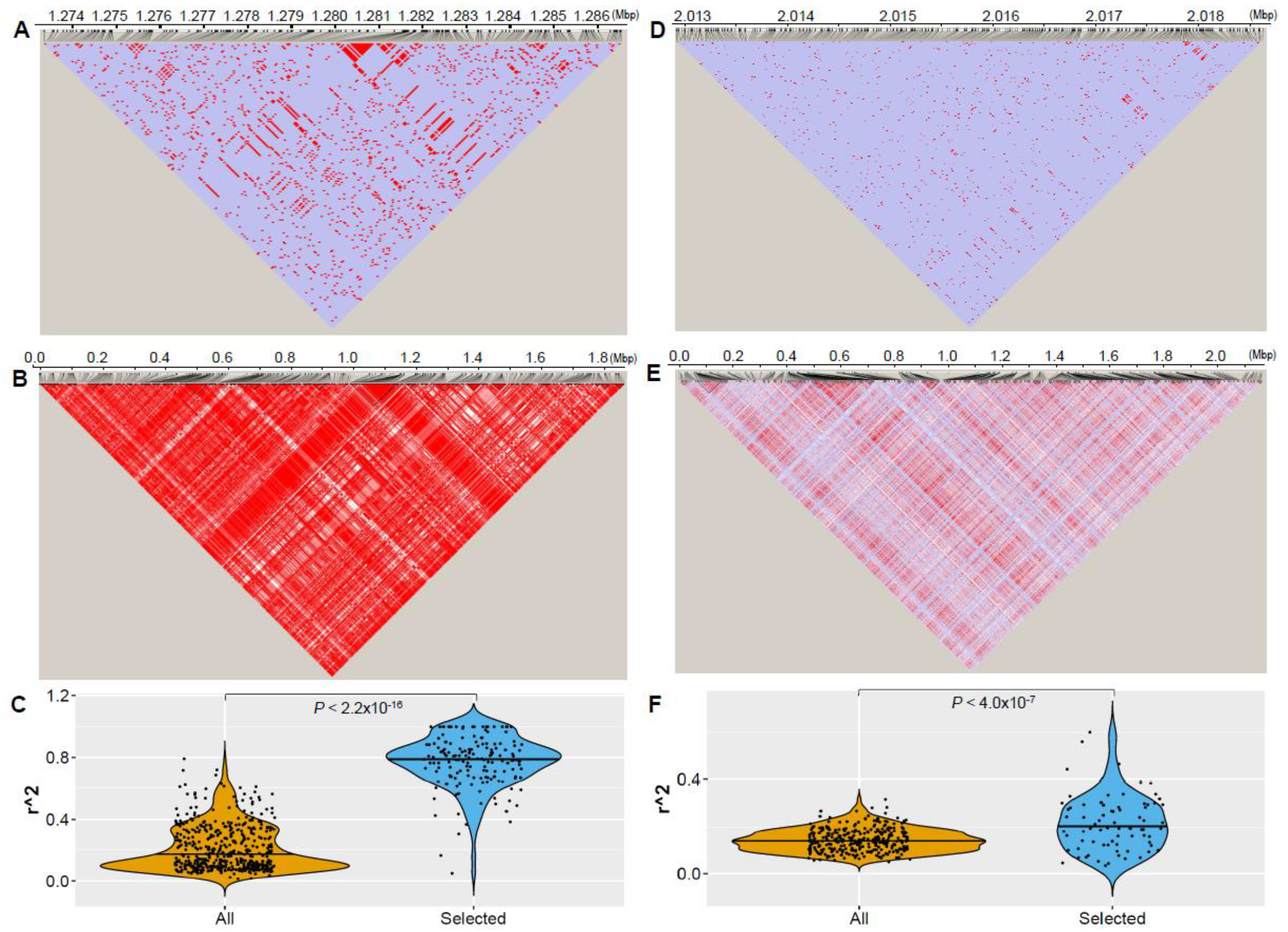
Comparison of the LD patterns for the SNPs before and after pre -selection for the genotype datasets of *S. pyogenes* (A,B,C) and *S. suis* (D,E,F). (A,D) The LD pattern of SNPs in the most significant cluster for all segregating SNPs from *S. pyogenes* and *S. suis*, respectively. (B,E) The LD pattern of the selected SNPs with phenotypic association in *S. pyogenes* and population differentiation in *S. suis*. (C,F) Distribution of the average level of inter-SNP LD in the clusters for all segregating SNPs and the selected subset of SNPs from *S. pyogenes* and *S. suis*, respectively. The LD pattern in (A) involves 1,014 SNPs located in the genomic region 1,273,267-1,286,739 of *S. pyogenes* AP53. The pattern in (B) involves the same set of SNPs as those used in Fig. 5E of Ref.Bao and includes 1,631 SNPs associated with acute rheumatic fever. The LD pattern in (D) involves 1,787 SNPs located in the genomic region 2,012,889-2,018,654 of *S. suis* BM407. The pattern in (E) includes 2,205 SNPs associated with population differentiation of *S. suis*. The LD patterns were generated by Haploview based on the pair-wise measure of the linkage disequilibrium D’ and log likelihood of odds ratio LOD. The different LD levels are indicated in color with red for the strongest LD (D’=1 and LOD > 2), pink for the intermediate LD (D’ < 1 and LOD > 2) in pink, white for the weak LD (D’ < 1 and LOD < 2) in white, and purple for uninformative (D’ = 1 and LOD < 2). The average inter-SNP LD (measured as correlation coefficient r^2^) was significantly increased for SNPs subject to pre-selection. The between-group difference was evaluated using Wilcoxon rank-sum test.

We carried out similar analysis for the genomic data of *S. suis* as that for *S. pyogene*. A total of 236,860 core SNPs were obtained across 208 non-redundant strains of *S. suis* and 349 clusters were identified using SweepCluster (*p*-value 0.05) (Figure S4A, **Additional file 2**: Table S6, S7 and S8). Without pre-selection of SNPs, we found that the clusters are densely distributed on the genome, implying that many of the clusters may contain false positive signals of selective sweep. Therefore, we selected the SNPs associated with differentiation of two subpopulations using the Chi-squared test (Figure S1C). A total of 2,205 SNPs satisfies the significance threshold (*p*-value 0.05) and were subject to cluster detection using SweepCluster (**Additional file 2**: Table S9). A total of 111 clusters were identified with significance (*p*-value 0.05) (Figure S4B and **Additional file 2**: Table S10). We examined the effect of SNP pre-selection by calculating the average inter-SNP LD within the clusters (Figure 4D,E,F). The results reveal a higher level of average LD in the clusters from the selected SNPs than that from the whole set of SNPs (*p*-value < 4.0 × 10^−7^), reiterating the efficiency of our strategy for identification of signals of sweep regions.

### Application in empirical datasets of *V. cyclitrophicus*

We benchmark our method using the dataset in the Ref. (9), the only available study of SNP clusters under gene-specific sweep in bacteria. We processed the genomic data from the 20 strains of *V. cyclitrophicus* (13 L strains and 7 S strains) to obtain ecoSNPs associated with ecological differentiation between the L and S population (**Additional file 3**: Table S11 and S12). Cluster detection is subsequently performed to the ecoSNPs using SweepCluster and 11 significant clusters were identified (Figure 5, Table 1 and Table S13). We validated our results by comparing with all eleven but two clusters reported in the Ref. (9). We excluded cluster2 annotated as “Conserved protein” of which the equivalent gene in our reference cannot be precisely located, and cluster4 which contains flexible genes without falling into the core genome. Among the remaining nine clusters, seven were recovered by our method corresponding to a concordance rate of 78%. It is noted that cluster5 was not recovered because it does not contain non-synonymous or upstream regulatory mutations, reflecting different clustering strategies of the two studies. It is noticeable that we also identified with high significance two novel clusters cluster12 and cluster13 containing 6 and 36 SNPs, respectively (Table 1 and Table S13).

**Table 1.** List of gene clusters identified by SweepCluster for ecoSNPs in *V. cyclitrophicus*.

| Chr. | ClusterID<br>in Ref. | First SNP | Last SNP | Cluster<br>size | Cluster<br>length | *P*-value | Gene loci range |
| --- | --- | --- | --- | --- | --- | --- | --- |
| chr1 | 1 | 2,999,463 | 3,017,933 | 50 | 18471 | $< 10^{-8}$ | FAZ90_RS13520-FAZ90_RS13590 |
| chr1 | 3 | 3,022,425 | 3,038,568 | 53 | 16144 | $< 10^{-8}$ | FAZ90_RS13690-FAZ90_RS13840 |
| chr1 | 3 | 3,044,230 | 3,045,684 | 63 | 1455 | $< 10^{-8}$ | |
| chr1 | 3 | 3,052,101 | 3,059,345 | 158 | 7245 | $< 10^{-8}$ | |
| chr2 | **12** | 307,956 | 307,979 | 6 | 24 | $< 10^{-8}$ | FAZ90_RS16555 |
| chr2 | 11 | 353,294 | 354,071 | 27 | 778 | $< 10^{-8}$ | FAZ90_RS16800 |
| chr2 | 10 | 533,257 | 533,284 | 6 | 28 | $< 10^{-8}$ | FAZ90_RS17530 |
| chr2 | 8 | 744,397 | 745,417 | 35 | 1021 | $< 10^{-8}$ | FAZ90_RS18465-FAZ90_RS18470 |
| chr2 | **13** | 1,514,577 | 1,525,409 | 36 | 10833 | $< 10^{-8}$ | FAZ90_RS21880-FAZ90_RS21930 |
| chr2 | 7 | 1,642,124 | 1,642,167 | 2 | 44 | $1.2 \times 10^{-5}$ | FAZ90_RS22415 |
| chr2 | 6 | 1,685,459 | 1,686,320 | 103 | 862 | $< 10^{-8}$ | FAZ90_RS22615 |

**Fig. 5.**
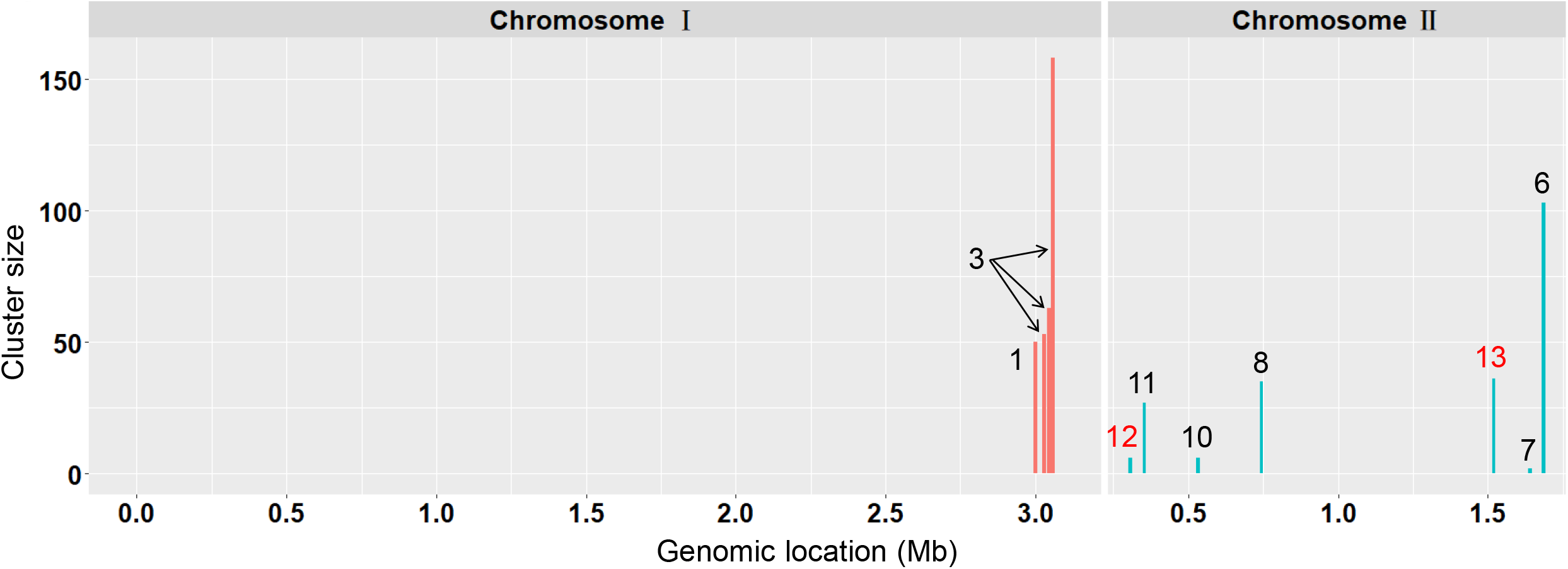
SNP clusters with signatures of selective sweep identified by SweepCluster for ecoSNPs of *V. cyclitrophicus*. The clusters are represented as colored bars with the bar height indicating the number of ecoSNPs in the clusters. Previously reported clusters in Ref.Shapiro recovered by SweepCluster are indicated in black numbering from 1 to 11 and those new clusters identified by SweepCluster indicated in red from 12 to 13.

In summary, the cluster comparison shows that the differences in the identified clusters between our results and those in the Ref. (9) are mainly due to distinct clustering methods and reference genomes used in the two studies. The current study used the strain of *V. cyclitrophicus* ECSMB14105, the only strain of this bacterium with complete genome assembly, while the study of (9) took an alternative but closely related species *V. splendidus* (12B01) as the reference. Therefore, the concordance rate between the two studies should have been underestimated.

### Application in empirical human genotype datasets

Though SweepCluster is specifically developed for prokaryotic data of dense genotypes, it will be helpful to test whether it is also applicable to eukaryotic data. We examined three well-characterized gene regions (LCT, EDAR, and PCDH15) under selective sweep in pairwise populations of EUR, AFR, and EAS from the human 1000 Genomes Project genotype datasets (30). We at first performed SNP pre-selection based on the population differentiation *F*st at a series of cutoff values, and then applied SweepCluster to each dataset of selected SNPs to search for gene regions under potential selective sweep (**Additional file 4-7**). At the threshold of *F*st = 0.4, all three gene loci were recovered as significant regions under selective sweep (Figure 6). The LCT gene, encoding lactase, was previously shown to be associated with lactase persistence in European populations and the region around it has been acknowledged as the target for strong selective sweep (19, 20, 34). In our cluster detection, the LCT locus along with the flanking gene regions (R3HDM1, UBXN4, and MCM6) forms a cluster of 57 variants spanning 235.6 kb with significance (*p*-value = 5.7 × 10^−6^), consistent with the strong positive selection. The gene EDAR is involved in ectodermal development and the missense mutation V370A showed evidences for association with hair thickness in East Asians (35,36). The region around EDAR has been identified to be the locus undergoing strong selective sweep (19, 36, 37). We localized the EDAR-centered region (GCC2, LIMS1 and EDAR) of 132 variants (including V370A) spanning 145.8 kb with significance (*p*-value < 10^−8^), implying strong selection signals. The gene PCDH15 encodes protocadherin and previous studies showed evidences of positive selection in East Asian populations (37, 38). We recovered the PCDH15 locus as a highly significant sweep region consisting of more than 300 variants spanning 369.5 kb (*p*-value < 10^−8^), indicating a strong signature of selective sweep.

**Fig. 6.**
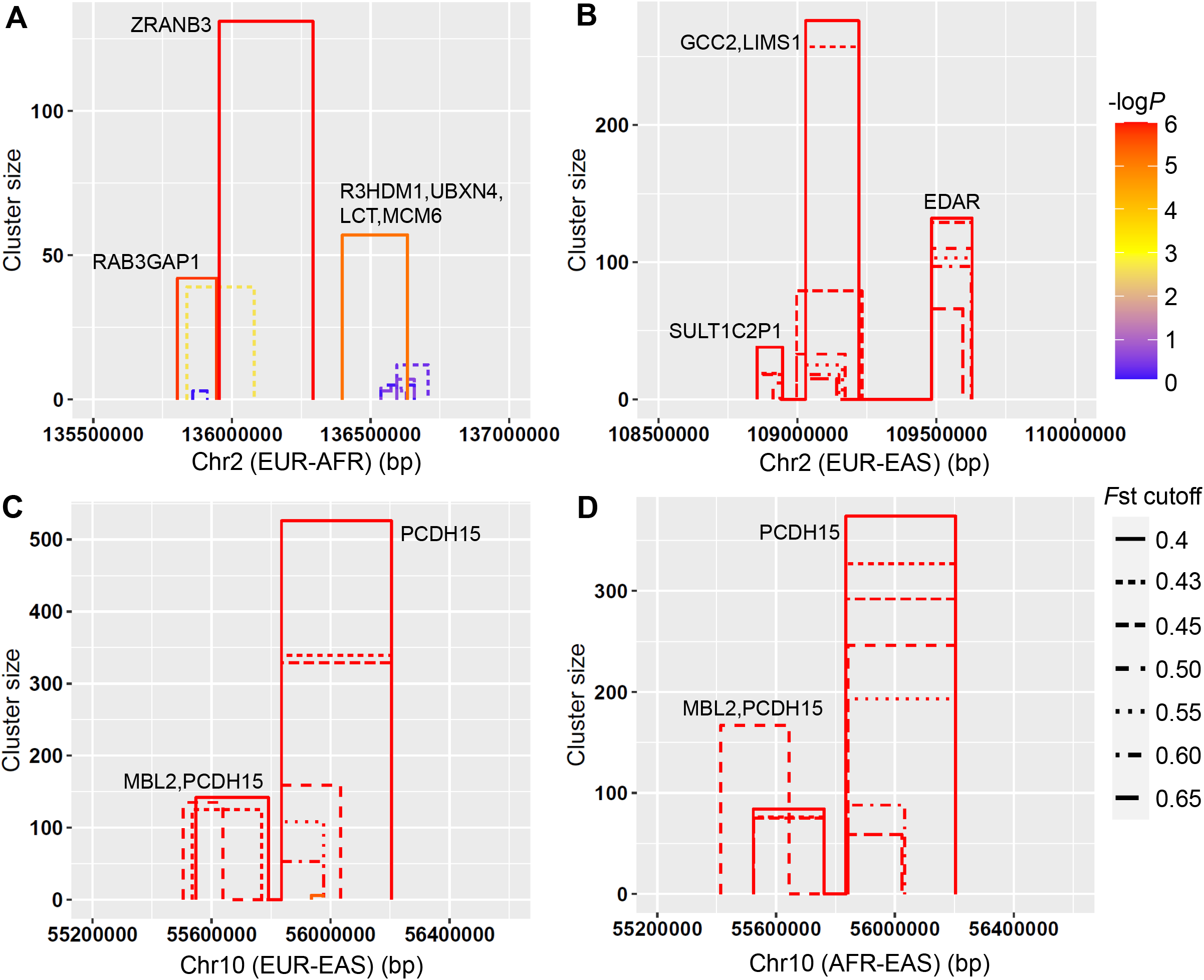
Sweep regions recovered by SweepCluster at three gene loci in the human 1000 Genomes Project genotype datasets. (A) LCT, (B) EDAR, and (C,D) PCDH15. A series of SNP selection criteria of *F*st were used to obtain distinct genotype datasets. The sweep regions are represented as colored bars with the bar height indicating the number of SNPs in the region (or cluster size) and the bar width indicating the spanning length. The significance is shown in -log10 (*p*-value) indicated in gradient colors.

Our results show that the size and significance of the sweep regions depend on the SNP selection threshold of *F*st, but the detection efficiency is robust for a wide range of *F*st. The signals of selective sweep emerge in all three gene regions at the threshold of *F*st = 0.4, and persist until *F*st ≥ 0.7. Above this threshold, the sweep signals in all three genes disappear. It is because a low number of mutations remain at the high level of *F*st and are sparsely distributed across the chromosome, making spatial clustering of the mutations inaccessible. We conclude that SweepCluster is also capable of detecting sweep regions for eukaryotic genotype data and the detection efficiency is robust to the SNP selection criteria.

### Optimization of parameters

The performance evaluation based on simulation data showed that the performance of the clustering algorithm in SweepCluster is closely related with the sweep length. Therefore, proper estimation of sweep lengths is critical for confident inference of selective sweep regions. Unfortunately, in many cases, it is not straightforward to derive the value of sweep lengths from genotype data. Therefore, we provided in the package a simulation script “sweep_lg_simulation.sh” to search for the optimal estimation of the sweep length for a specific genotype dataset. It is particularly suitable for prokaryotic data because the prokaryotes use gene conversion as the main vehicle for introducing selective sweeps and the sweeps are generally uniform in size (21).

We did the simulation by calculating the number of sweep regions inferred by SweepCluster at a series of values of sweep lengths and then fitting a non-linear model for the relationship between the number of sweep regions and sweep lengths. The optimal estimation of sweep lengths is determined by the point of maximum curvature in the fitting model. Here we present the simulation results for the three real datasets of *S. pyogenes, S. suis*, and *V. cyclitrophicus*, respectively (Figure 7). It is shown that all three datasets have the maximum curvatures at the sweep length of ∼2000 bp (1638 bp for *S. pyogenes*, 1500 bp for *S. suis*, and 2157/1989 bp for the two chromosomes of *V. cyclitrophicus*). It is consistent with our previous estimation of 1,789 bp for *S. pyogenes* using the alternative tool ClonalFrame (10, 39).

**Fig. 7.**
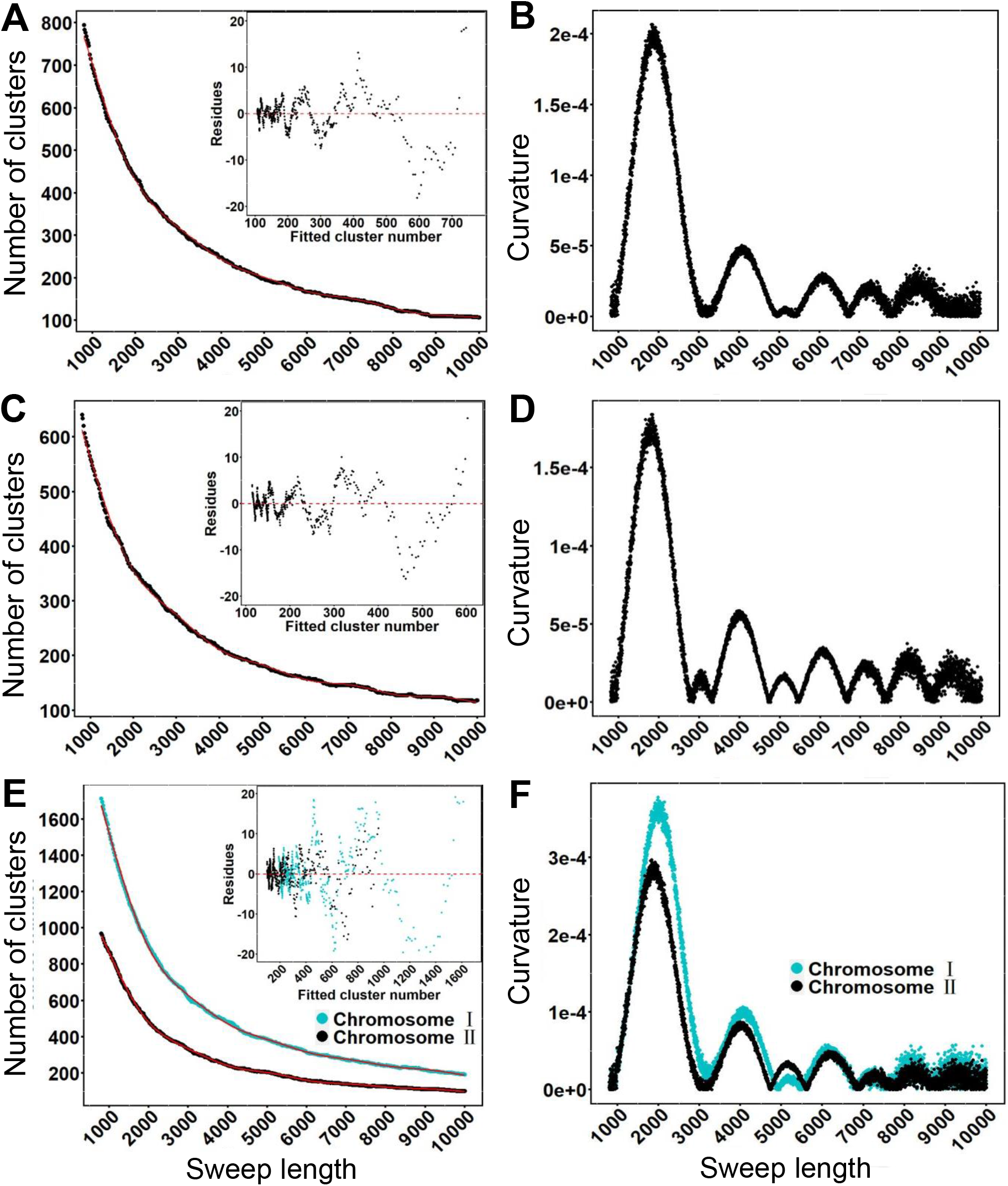
Parameter optimization of sweep lengths based on non -linear fitting and maximum curvature in three prokaryotic datasets. (A,B) *S. pyogenes*. (C,D) *S. suis*. (E,F) *V. cyclitrophicus*. The relationship between the number of sweep regions and sweep lengths was fit using generalized additive models (red lines) and the curvature for each fitting curve was calculated using the formula (4).

## Discussion

We have proposed a gene-centric spatial clustering approach to identify gene-specific sweeps in bacterial polymorphism data. It targets for the mutation sites complying with specific genetic properties of selective sweeps and captures the regions with unusual clustering patterns of those mutations differing from that of a neutral expectation. Based on the known genetic properties of gene-specific sweeps, the target mutations are usually obtained by selecting those with elevated population differentiation, reduced within-population diversity, heightened linkage disequilibrium, or significant phenotype association. The selected subsets of mutations are subject to clustering. Therefore, our approach for inferring sweep regions employs two layer of information, *i*.*e*., genetic signatures and spatial distribution patterns of mutations under gene-specific sweeps in comparison with current methods focusing only on one layer of information in the genotype data (11-14, 16, 19, 20).

The purpose of the procedure of selecting target mutations of particular genetic signatures prior to clustering is to remove the spurious or uninformative signals and perform spatial clustering only for the mutations related with selective sweep. The impact of mutation selection was dramatic in our two example datasets from the bacteria *S. pyogenes* and *S. suis*. The level of linkage disequilibrium between SNPs, as a signature of selective is significantly increased by selecting those mutations associated with disease phenotypes or population differentiation. The ultimate datasets upon prior selection are more sensitive to the statistical test under the neutral model of spatial distribution of mutations, making it more efficient to identify gene regions under selective sweeps. Using the only available dataset of gene-specific sweeps in bacteria (9), we validated our method yielding a concordance rate of 78% for the detected clusters even with distinct clustering strategies and reference genomes in the two studies.

Our approach is specifically designed for prokaryotic data of dense genotypes such that the mutations of particular genetic properties can be exhaustively obtained and the distribution of those mutations can be statistically distinguished from the null model. However, the testing showed that our method also performs well for eukaryotic data of sparse genotypes. We recovered the well-characterized gene regions (LCT, EDAR, and PCDH15) under selective sweeps in the 1000 Genomes Project genotype datasets. The signals of selective sweeps in the three gene loci persist for a wide range of mutation selection criteria, suggesting the robustness of our method on identifying sweep regions in sparse genotype data. Moreover, the spatial-aware strategy of the clustering makes the resolution of detected sweep regions narrowed down to single nucleotides facilitating identifying relatively old sweeps of low numbers of selected sites.

There are some limitations of our approach. It cannot distinguish explicitly between hard sweeps and soft sweeps, or recent sweeps and older sweeps because mixed sites of varying strength of selection are treated as a whole for statistical tests. Our method does not deal with the confounding effects of background selection, as the signatures of background selection are very similar to the real selection and it has been a challenge to confidently classify the background selection for many alternative approaches.

## Conclusion

We proposed a novel gene-centric approach for identifying gene-specific sweeps implemented in the Python tool SweepCluster. It performs spatial clustering of polymorphisms to infer the regions with signatures of gene-specific sweeps by employing two layers of information, *i*.*e*., genetic properties and spatial distribution models of the polymorphisms. It is specifically developed for prokaryotic data of dense genotypes and exhibit efficiency and robustness in detecting sweep regions in the validation datasets. It also performs well for eukaryotic data in a wide dynamic range of parameters of genetic properties. We expect that our new method will be valuable for detecting gene-specific sweeps in diverse genotype data and provide novel insights on evolutionary selection.

## Declarations

### Availability of data and materials

All data generated during this study is included in this article and its supplementary. The computational tool we developed “SweepCluster” is accessible at github: https://github.com/BaoCodeLab/SweepCluster.

## Acknowledgements

Not applicable.

## Figure legends

**Fig. S1.**
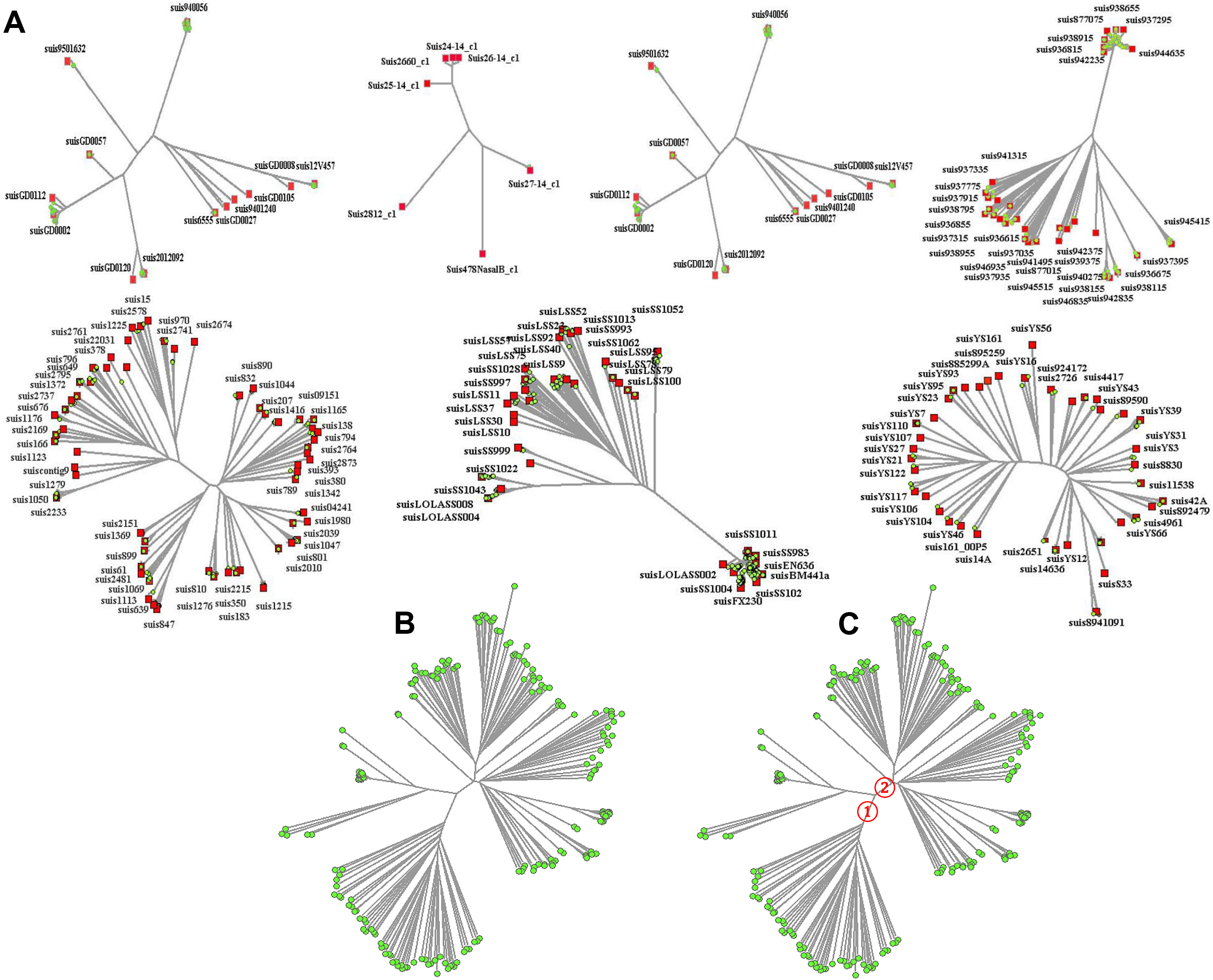
The phylogenetic trees for selection of non -redundant strains. (A) The trees for each of the seven groups of strains. The grouping was based on the submission institutions. The strains selected for downstream analysis in each group are indicated in red square. (B) The final phylogenetic tree for 208 selected non-redundant genomes. (C) The two subpopulations used for identification of SNPs associated with population differentiation are indicated with numbers.

**Fig. S2.**
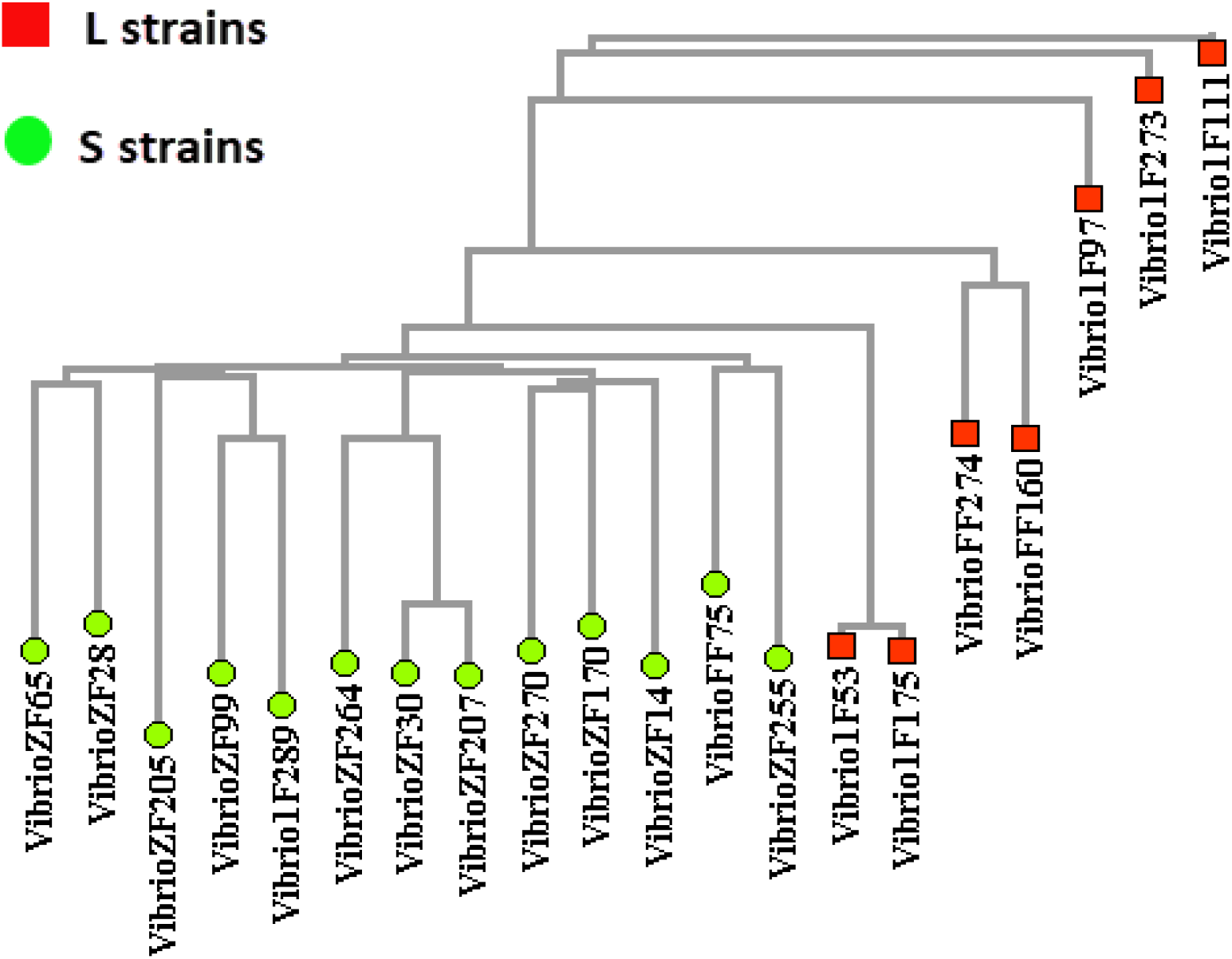
Phylogenetic tree of the 20 strains of *V. cyclitrophicus*. The ecological partition of the strains is indicated in color for thirteen S strains and seven L strains.

**Fig. S3.**
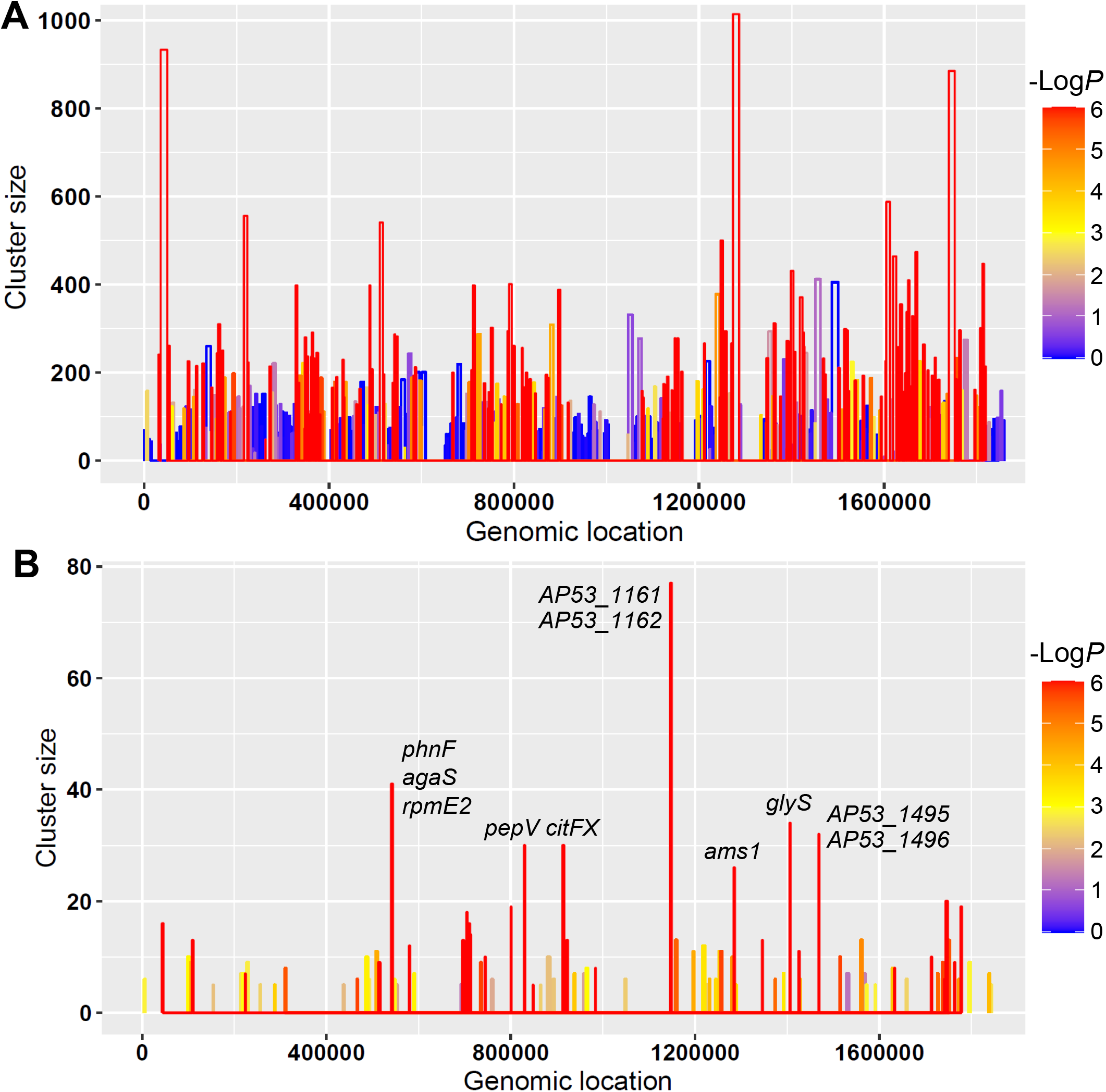
SNP clusters with signatures of selective sweep identified by SweepCluster for *S. pyogenes* genotype datasets. (A) The clusters detected from all segregating SNPs in the core genome. (B) The clusters detected from 1,631 selected SNPs with phenotypic association. The clusters are represented as colored bars with the bar height indicating the cluster size (the number of SNPs in the clusters) and the bar width indicating the spanning length. The significance of the clustering evaluated with -log10 (*p*-value) is indicated in gradient colors. The gene loci in the top clusters are shown.

**Fig. S4.**
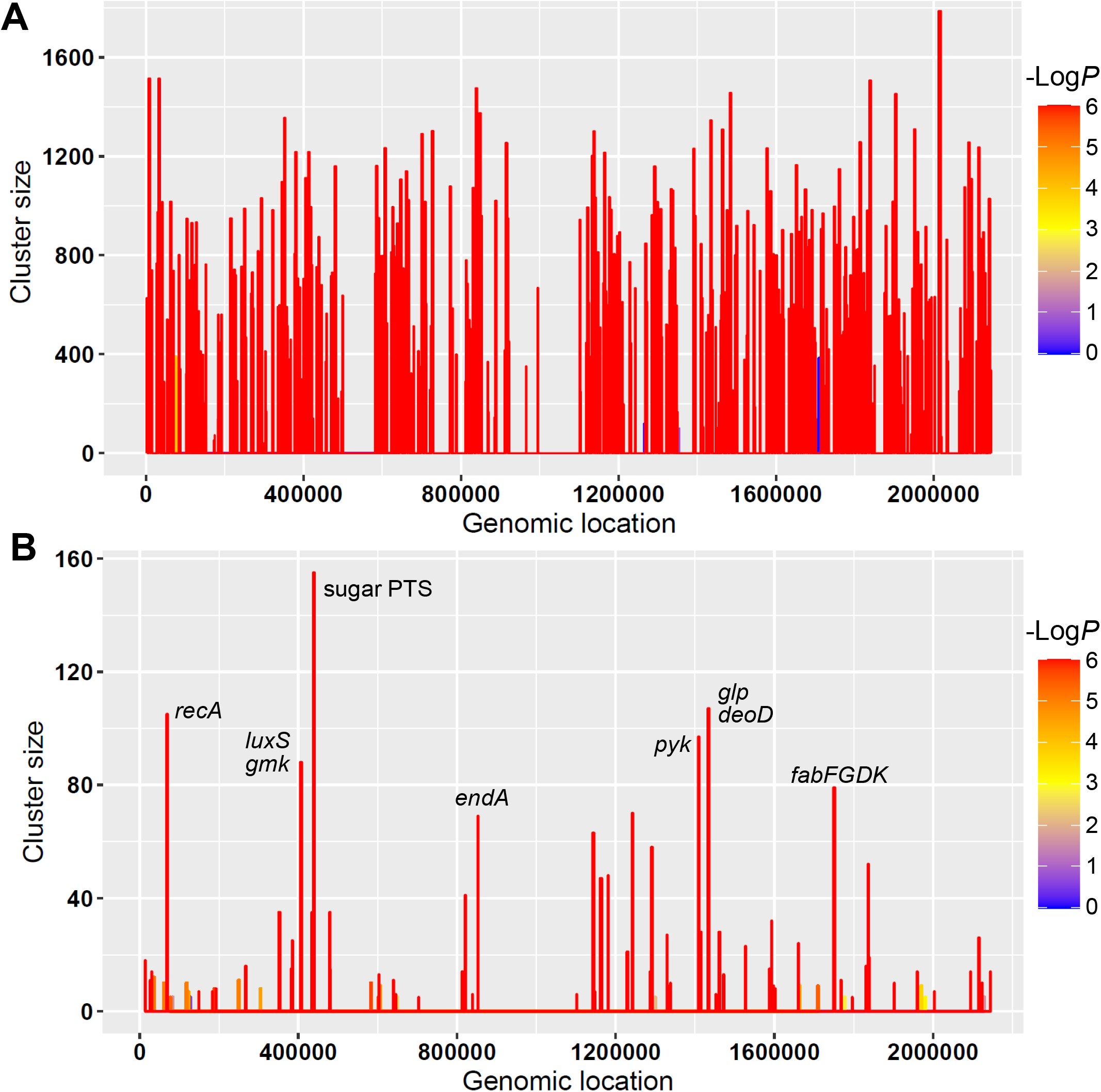
SNP clusters with signatures of selective sweep identified by SweepCluster for *S. suis* genotype datasets. (A) The clusters detected from all segregating SNPs in the core genome of *S. suis*. (B) The clusters detected from 2,205 selected SNPs associated with population differentiation (Chi-squared test *p*-value≤0.05). The clusters are represented as colored bars with the bar height indicating the cluster size (the number of SNPs) and the bar width indicating the spanning length. The significance of the clustering evaluated with -log10 (*p*-value) is indicated in gradient colors. The gene loci in the top clusters are shown.

